# Machine Learning Identifies Common Risk Variants and Implicates Abnormal Vision Physiology in ASD

**DOI:** 10.1101/2025.11.11.687806

**Authors:** Alona Rabner, Assaf Avrahami, Abraham Meidan, Ilya Vorobyov, Yael Mandel-Gutfreund, Guy Horev

## Abstract

Genomic technology advancements have facilitated associations between genetic variants and disease risk. Rare deleterious variants can independently induce disease, while common variants collectively enhance susceptibility with minimal individual effects. The nuanced nature of common variant consequences attenuates their identification. Autism spectrum disorders (ASD) are hereditary neurodevelopmental disorders. Atypical eye-gaze responses frequently occur in ASD; however, this phenotype has been overlooked in genetic studies. Using WizWhy, an interpretable machine learning tool, to analyze the Autism Sequencing Consortium (ASC) data, 210 common variants across 177 genes were associated with ASD risk. This association is supported by significant overlap with the SFARI gene database and relevant gene ontologies. Individuals with a higher variant burden are at increased ASD risk. Notably, 52 genes were linked to abnormal eye physiology, underscoring the role of this pathway in ASD etiology. Focusing on specific hub genes as potential pharmacological targets may benefit patients with ASD.

## Introduction

Autism spectrum disorders (ASD) are a group of neurodevelopmental conditions characterized by early onset impairments in social communication and interaction and restricted or repetitive behaviors(1). In 1977, Folstein and Rutter established a strong genetic contribution to ASD(2) and motivated molecular genetic investigations of ASD(3). A highly prominent deficit in social functioning, atypical social visual engagement, is observable within the first months of life in infants who are later diagnosed with autism(4), continues through later life(5), and is highly heritable(6).

Recent advances in genomic methodologies have enabled large cohort studies involving thousands of subjects at the genome and exome scales(7). These studies found that de novo and rare variants, mainly structural and protein-disrupting variants, are liable to increase individual ASD risk(8–14). Together, variants in hundreds of genes can explain 10–40% of ASD cases. These findings indicate that ASD is highly heterogeneous at a genetic level. High heterogeneity limits the predictive power of these variants and, hence, the ability to translate rare variants for use in diagnosis or medical interventions.

In contrast to rare variant research, studies of common variants address the question of whether there is a contribution to the ASD phenotype from variants with small effects. These studies assumed an additive effect of multiple variants harbored by one individual(15). Several reports have provided evidence for this model, and estimates indicate that common variants explain more than 50% of heritability(16,17). A meta-analysis of 18,381 affected individuals and 27,969 controls led to the identification of five genome-wide significant loci. Further work, however, is required to uncover the likely causal genes within these loci and how gene variation leads to the disorder(16).

In addition to the challenge of identifying common causal variants, association studies assume that a trait is under neutral selection. They also require that the cases and controls are ethnically matched, but not close relatives, and that the underlying liability follows a Gaussian distribution(18). These assumptions are not fulfilled in ASD; for example, it is not a neutral trait. Hence, A2DS, a non-parametric method for identifying common risk variants in cohorts of discordant sibling pairs, was developed(18). Notwithstanding, case–control design is common in genetic studies, and a large amount of data has already been collected on ASD and other neurodevelopmental disorders. Therefore, we aimed to identify common variants in ASD using an interpretable machine learning approach that does not rely on the aforementioned assumptions within a case–control study.

In this study, we used WizWhy(19), a machine learning software program designed to associate independent variables (e.g., genetic variants) with the dependent variable (e.g., phenotype). These associations are referred to as rules. WizWhy works by revealing all if– then rules that associate combinations of the independent variables with the dependent variable and meet user-determined thresholds for rule confidence (minimum probability) and support (minimum number of cases). The software uses these rules to predict the subject diagnosis in a test dataset. A key advantage of using WizWhy is that the rules provide explanations for the predictions and can be used to build hypotheses that explain the data.

We trained WizWhy to generate rules that are based on common variants and distinguish between affected and control individuals. We then extracted rules contributing to the software’s ability to distinguish between the two groups. The rules provided were heavily biased toward identifying affected individuals, as opposed to a randomized dataset that provided rules matching the proportions of affected and non-affected individuals. Noting that due to a much higher number of independent factors than the number of subjects in the study, many rules can be false positive, we discarded rules that did not hold in an independent cohort of controls. This process identified 536 variant pairs in 177 genes that distinguished reliably between affected and unaffected individuals. Of the 177 genes, 18 are listed in the SFARI(20) gene database, a significant overlap indicating that common and rare variants leading to ASD converge on similar pathways. We provide evidence for an additive effect of the variants: the more risk variants a subject harbors, the higher the chances of being diagnosed with ASD. Interestingly, enrichment analysis highlighted an abnormal eye physiology pathway. Finally, hub genes that appeared in a higher proportion of variant pairs were identified. We propose these hubs as targets for drug development because their variants are common to many affected individuals. Collectively, these results provide new opportunities for ASD diagnosis and treatment.

## Results

### Data preparation

We explored exome sequencing data from 8,061 individuals—3,585 diagnosed with autism and 4,476 controls (see Methods)—from the Autism Sequencing Consortium (ASC)^21^. The analysis involved variants associated with changes in protein sequence including protein-truncating, frameshifts, and missense variants. To reduce the number of potential combinations of independent variables and enable us to focus on abundant variants, we implemented a two-step approach. First, we quantified the frequency of each variant across the entire cohort. Subsequently, for each participant, in our dataset we included only the most common variant within each gene that the individual carried (see Methods). This procedure allowed us to prioritize the most frequently occurring single nucleotide polymorphisms (SNPs) while maintaining a comprehensive view of every individual’s genetic profile. This method resulted in 18,497 genes harboring 655,780 variants. To construct a control dataset, we randomized the order of variants in each gene to generate a dataset with similar variant distribution but without association between genotype and ASD. The data were further divided randomly into three non-overlapping sets of subjects. A training set, consisting of 70% of the study subjects, and pruning and test sets, each comprising 15% of the subjects. This procedure was repeated five times with each dataset, to produce five training–pruning– test cross-validations of the original and randomized sets.

### WizWhy reveals meaningful associations between ASD and common variants

In construction of the prediction model, WizWhy’s objective is to minimize the total cost of errors. The user specifies the cost of a false negative and the cost of a false positive. In the present study, these error costs were inversely proportional to the frequencies of the dependent variable’s values. During the process of generating predictions, WizWhy performs the computation of two key components: (A) the minimum total cost of errors when making predictions based on the frequencies of the dependent variable and the associated costs of errors, and (B) the total cost of errors associated with the predictions generated by the model. The ratio between A and B represents the improvement factor. Training and testing were conducted using both the original and randomized data. Training on the original dataset resulted in an average improvement factor of 1.14 [1.1-1.17] in predicting whether an individual is affected or not. Conversely, the utilization of the randomized dataset yielded an average improvement factor of 0.95 [0.92-0.99]. This result indicates that when using the original dataset, WizWhy produces predictions superior to random predictions as to whether an individual belongs to the case or control group, but not when utilizing the randomized data. This finding suggests that the genetic variants in our dataset contain relevant information pertaining to ASD. Examination of the rules revealed that WizWhy identified between 1,643,223-1,957,185 rules in each cross-validation of the original data, compared to 67,000-91,000 rules when using the randomized data. For each WizWhy run, the strongest if-then rules are reported up to a limit of 100,000 rules. The rules generated for the randomized data are false positives, indicating that false positives are also present within rules generated with the original (non-randomized) data. The observation that the number of rules identified in the original data is more than 10 times higher than in the randomized data, however, indicates the discovery of true positive rules. Notably, 80-90% of the rules generated by training with the original data were biased toward affected outcomes, while rules derived from the randomized dataset were biased toward non-affected outcomes. Only 30-35% of the rules in the randomized sets indicated affected outcome, in accordance with the 45% proportion of affected in the training sets. The strong bias toward prediction of affected individuals in the original dataset suggests that SNPs are associated with deleterious effects rather than with protective effects in control individuals. In summary, WizWhy revealed significant associations between common variants and ASD risk in the original data. Thus, to further identify variants that increase the risk of ASD diagnosis, it is necessary to differentiate valid rules from false positives.

### Verification of variant pairs in a distinct unaffected cohort

The if-then rules elucidating WizWhy’s predictions encompassed a total of 100,000 rules within each cross-validation round, summing to 415,246 rules predicting affected outcomes. A substantial proportion of these rules were observed across multiple iterations resulting in 190,311 unique rules (Supplementary table 1). To differentiate genuine signals from overfitted information, a decision was made to evaluate these rules on an independent cohort, specifically the control group derived from the Swedish Schizophrenia (SCZ) Population-Based Case-Control Exome Sequencing study (phs000473.v2.p2), comprising 5,890 participants. Given that all the rules tested involved two variants in distinct genes, the rationale underlying this approach was that variant pairs absent in both control cohorts (ASD and SCZ studies) are more likely to represent accurate rules, thus reducing the proportion of false positive rules. Consequently, the expected occurrence of each pair among individuals was computed based on their frequency in the SCZ control group dataset, followed by the application of the Fischer Test to identify pairs significantly less prevalent in the SCZ control cohort than would be expected by chance. The focus was on selecting variant pairs where each variant is common, yet their joint occurrence is rare among a non-affected cohort, indicative of a process of purifying selection acting on these specific pairs. Following adjustment for multiple comparisons (adjusted p-value < 0.05, False Discovery Rate (FDR)^22^), a total of 585 distinct pairs involving 193 unique genes and 221 SNPs were identified. Among these genes, 18 belonged to three clusters of the protocadherin family, located on chromosome 5 and the same SNP was mapped to different members of the same cluster. To prevent an overrepresentation of these related family members in the dataset that occurs due to mapping problem, a single representative protocadherin was selected from each cluster, resulting in a refined dataset of 536 distinct pairs and 177 unique genes (Supplementary table 2).

### Additive effect of the discovered variants on ASD risk

After verifying 210 common variants in 177 genes, we examined their distribution within this cohort. Common variants have the potential to impact individuals cumulatively, through the aggregation of numerous variants, each exerting a minor effect that contributes to the development of ASD. Alternatively, interactions between variants may also play a significant role, given that individual variants with modest effects combine synergistically. The result is the manifestation of an effect that surpasses the sum of the effects of each individual variant.

The heatmap presented in Figure 1A provides evidence for an additive effect of the variants. By arranging the participants in ascending order of the number of variants they possess, a substantial proportion of affected individuals are situated in the lower section of the heatmap. Particularly, individuals harboring 32 variants or more have all been diagnosed with ASD. Looking at the individuals in the cohort harboring at least one of the 536 rules shows a distinct distribution of the 210 risk variants in affected and control subjects (Fig. 1B). While control individuals have less than 10 risk variants on average, affected individuals have more than 30 risk variants on average (p-value< 2.2e-16, t-test). A generalized linear model indicated that the probability of being affected increases with the number of risk variants harbored by a subject (Fig. 1C). This analysis strongly suggests that an elevated number of variants within an individual escalates their risk of ASD.

**Figure 1.**
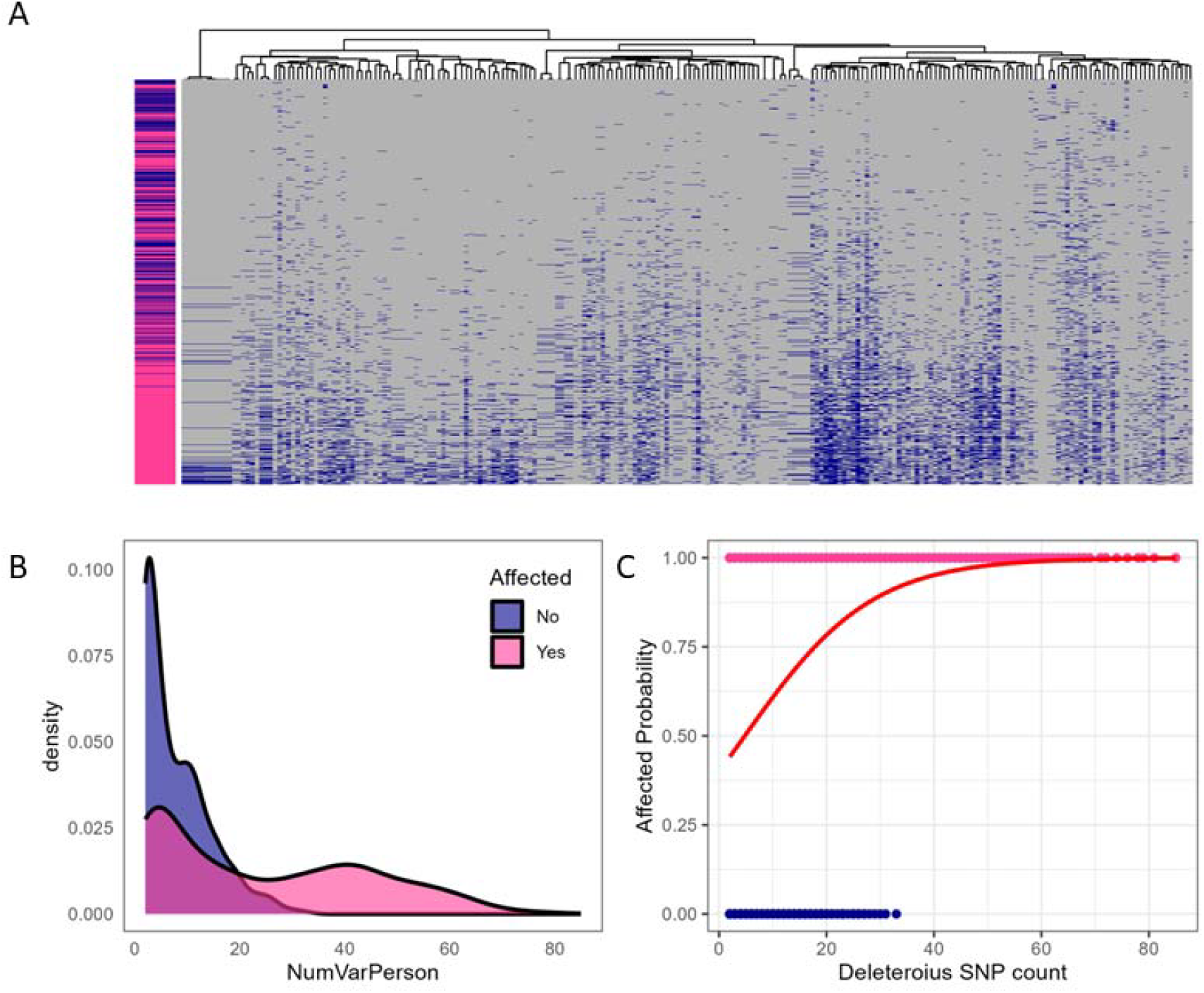
Additive effect of common SNPs on ASD risk. A. A heatmap depicts the distribution of SNPs (columns) among subjects (rows). Subjects are ordered in ascending variant burden (lowest burden at the top, highest at the bottom). The side bar indicates the affected status of each subject (blue = not affected, violet-red = affected). The plot shows that individuals with a higher variant burden (located towards the bottom of the graph) are more frequently affected. B. Density plot showing significantly higher (p-value< 2.2e-16, t-test) deleterious SNP count in affected subjects. C. Logistic regression shows that the probability of being affected increases as the deleterious SNP count grows (p-value< 2e-16, generalized linear model).

### Common and rare variants leading to ASD converge on similar pathways

Thus far, most ASD risk variants were discovered through rare variant investigations. A key question arises as to whether the potential impact of common variants within the same gene pool is the same as rare variants, or whether the former influence distinct genes. To address this question, we assessed the intersection of the 177 genes identified in this study with the SFARI Gene database, which focuses on genes linked to autism susceptibility through the identification of rare variants. Markedly, 18 genes were found to coincide with the 1136 genes in SFARI, indicating a significant overlap (p=0.025, hypergeometric test, Fig. 2A, Supplementary table 2). This overlap implies a convergence of rare and common variants in ASD on similar pathways. Gene ontology enrichment analysis of the 177 genes using ShinyGO revealed significant enrichment for neuronal and brain development processes (Fig. 2B,C, Supplementary table 3). Hierarchical clustering was performed to determine the extent of overlap between the GO terms associated gene sets with Jaccard similarity. This divided the GO terms into two clusters. Cluster one involves five GO terms that include a small number of genes, each showing high fold change, but no overlap with other significant GO terms. Two of these pathways belong to the sarcomere organization processes. The other three are notch binding, high voltage-gated calcium channel activity and eukaryotic translation initiation factor 2. Cluster two involves four terms all belong to neuron projection and differentiation processes. All these ontologies were previously associated with ASD in studies of rare variants(27–30), supporting the inferences that common and rare variants converge on similar pathways in ASD.

**Figure 2.**
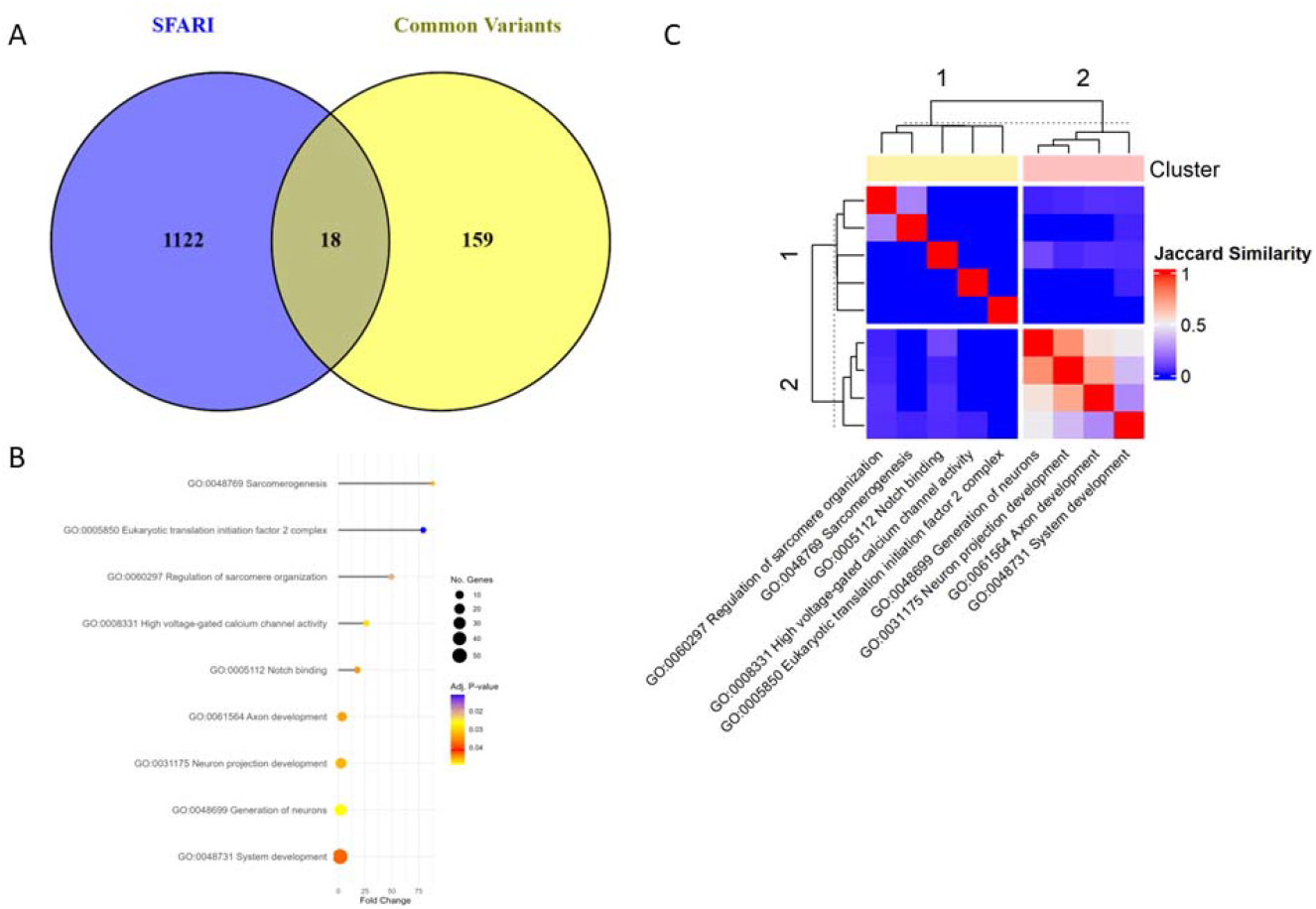
Common and rare variants leading to ASD converge on similar pathways. A. Venn diagram showing significant overlap (p=0.025, hypergeometric test) with the SFARI database. B. Enrichment of GO term annotations. C. A heatmap representing a hierarchical clustering among the significant GO terms listed in B.

### Abnormal eye physiology implicated by common variants in ASD

To further our enrichment toward potential phenotypes affected by common variants, we focused the Human Phenotype Ontology (HPO) terms (31) in the ShinyGO enrichment analysis. This analysis resulted in 48 significant (adjusted p-value<0.05) terms (Fig.3A), many are ASD related phenotypes as detailed below. These 48 terms were clustered into 5 clusters based on the overlap of their gene sets (Fig.3 B). Clusters one, two and four show strong overlap in their gene content. Cluster 4, the largest cluster (24 terms), harbours two core ASD phenotypes: neurodevelopmental delay and abnormality of speech and vocalization, as well as many phenotypes that are common in ASD, seizure and motor delay to name two. Cluster two includes terms for infantile and pediatric onset that are core in ASD diagnosis. It also includes 64 genes in the term mendelian inheritance, meaning that variants in these genes are causal in hereditary diseases. Finally, cluster 1, includes terms that are all belong to abnormality of the eye phenotypes, in particular, abnormal eye movement phenotypes. The full list of significant phenotypes is provided in Supplementary table 4. This finding suggests that in some patients with ASD, abnormal eye movements may be causal in ASD.

**Figure 3.**
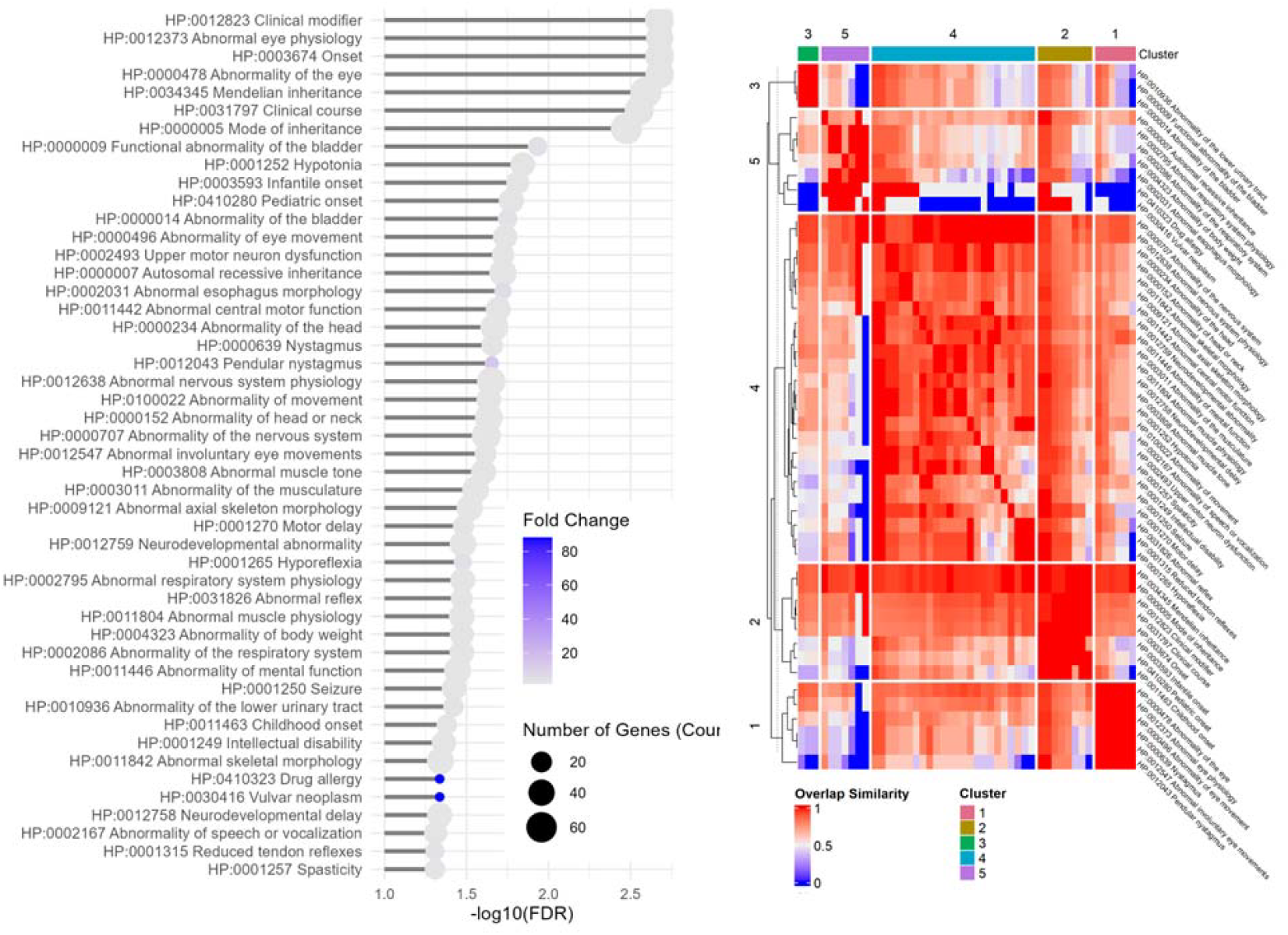
Abnormal eye physiology implicated by common variants in ASD. A. Enrichment of HPO phenotype annotations. B. Hierarchical tree of significantly enriched HPO terms.

ShinyGO enrichment using a subset of 49 genes overlapping with eye phenotypes highlights impairments in two molecular functions: low voltage-gated calcium channel activity and NOTCH1 regulation pathway (Supplementary table 5). Ten mammalian genes encode the α1 subunits of voltage-gated calcium channels. Three distinct SNPs in our analysis are harbored within three of these genes: *CACNA1G, CACNA1I*, and *CACNA1S*. The first two genes encode T-type channels involved in neuronal excitability and have been associated with ASD(32). *CACNA1S*, which encodes L-type channels, has also been implicated in ASD in rare variant studies(33). *NOTCH1* is a receptor protein that, when activated by binding to its ligands (such as Delta-like and jagged proteins), initiates a signaling cascade that influences cell fate decisions, differentiation, proliferation, and apoptosis(34). *NOTCH1* has been previously implicated in ASD^30^ but its *JAG1* and *JAG2* ligands have not been associated with ASD thus far. Interestingly, *JAG1* and *JAG2* exhibit calcium ion binding activity, suggesting potential crosstalk between *NOTCH1* and calcium channels.

### Gene connectivity in a network provides distinguishes between ontologies

A gene association network was constructed based on WizWhy-derived rules for disease prediction, where edges represent gene co-occurrence in the same predictive rule. The resulting network (177 nodes, 533 edges) provides insights into potential phenotypic outcomes. The number of connections a gene has in the network varies greatly. The most connected genes appeared in as many as 48 rules, while most genes appeared in 10 or less rules (Fig. 4). Three genes, MYLK3, CLIP1 and TNC stood out in the high number of rules co-occurrences, and are co-occur with each other. Network analysis showed that these 3 genes receive high scores in four complementary topological metrics (Supplementary table 6). Enrichment analysis using genes directly co-occur in rules with each of these 3 genes revealed genes that co-occur with MYLK3 and TNC are highly enriched for neuronal processes (Supplementary tables 7 and 8) while CLIP1 co-occurring genes did not show particular enrichment.

**Figure 4.**
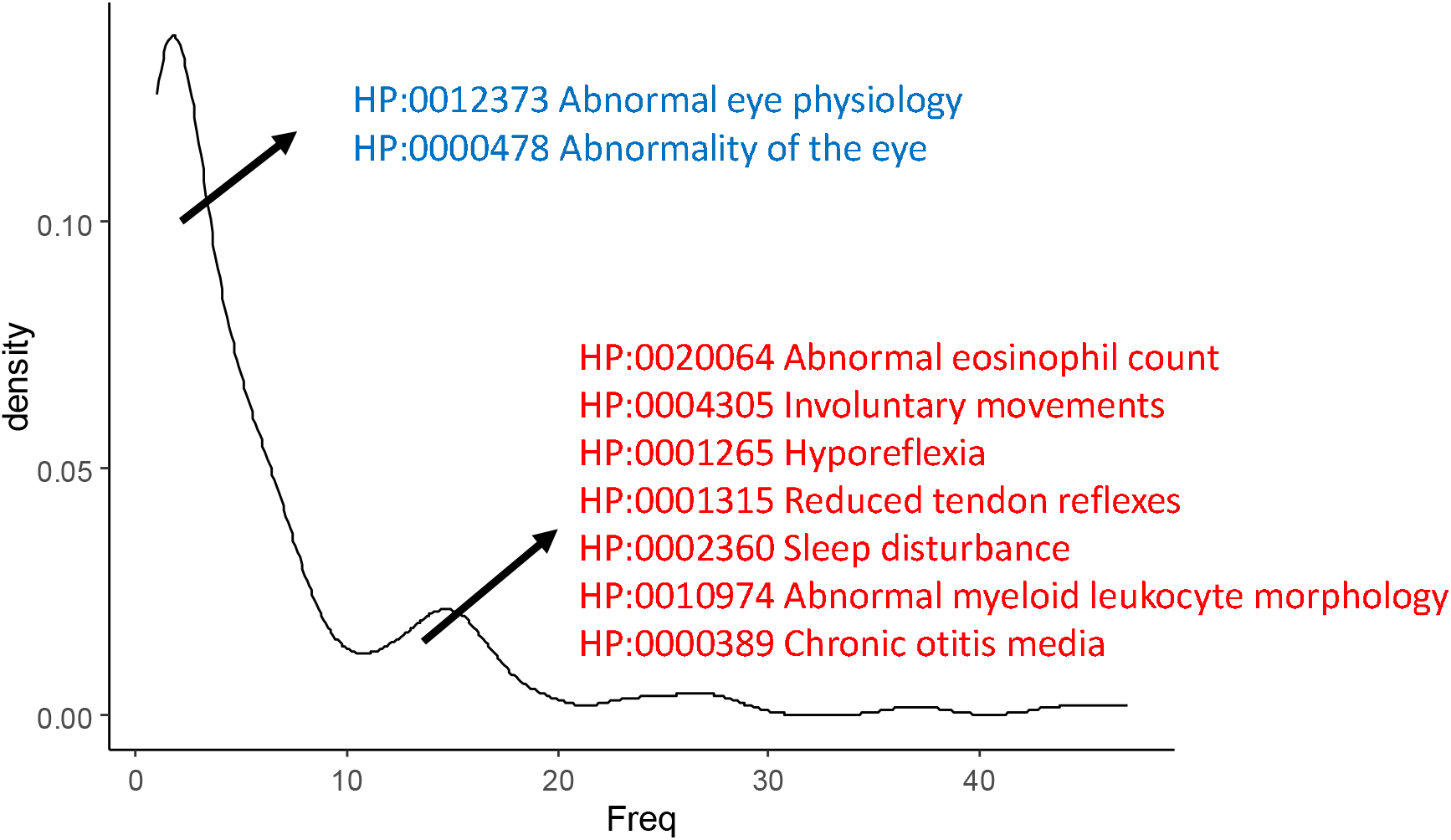
Gene connectivity determines distinct phenotypic enrichment – Density plot of the number of co-occurrences in rules. Genes with less than 11 connections are enriched for abnormality of the eye and genes with 11 connections or more are enriched for immune system, motor and behavioral phenotypes.

Enrichment analysis using the 144 low connectivity genes showed that these genes are responsible for the abnormality of the eye enrichment. In contrast, the 33 high connectivity genes showed significant enrichment for neuronal processes as well as enrichment for immune system processes such as B-cell activation and B-cell apoptosis. HPO enrichment of the high connectivity group showed enrichment to immune system phenotypes including allergy, asthma, and abnormal counts and morphologies of different immune system cell types. In addition, intellectual disability, hyporeflexia, and involuntary movements also appeared in this group.

Altogether, these results indicate that gene connectivity derived from the identified rules follows biological processes.

## Discussion

Common genetic variants play a significant role in influencing an individual’s susceptibility to various diseases. Associating common variants with disease risk, however, poses several challenges. One major obstacle is the need for large-scale genome-wide association studies (GWAS) with substantial sample sizes that enable researchers to detect the subtle effects of common variants^15^. In addition, the functional consequences of many identified variants remain unclear. Here, we demonstrate the power of machine learning based approach in uncovering subtle genetic associations that traditional methods may have overlooked.

First, we trained the machine to distinguish between affected and control individuals. We only included in the analysis the most common variant within each gene that an individual carried. This allowed us to reduce the analysis to a manageable size at the cost of ignoring many causal variants. This process resulted in 190,311 unique rules. The rule of thumb in machine learning is that to avoid significant overfitting, the number of records should be at least 10 times the number of independent variables. In our project, however, the number of independent variables was 18497 and the number of records in the training set was 5642. Therefore, overfitting (when predicting on the test dataset) was expected and indeed found.

Using the independent SCZ dataset as means for filtering false positives, we found valid rules. This approach tackles several challenges: First, by testing pairs of variants, we assess the additive effects of the variants and hence increase the power to detect small-effect variants. Testing in pairs also allows interactions to be detected, which may have a bigger effect than the additive effect of the two variants. Second, by testing the affected and non-affected cohorts separately, we avoid potential ancestry biases between affected and control cohorts that are inherent in case-control studies. Last, by constructing randomized datasets and performing machine learning on these datasets, we show that we do not issue meaningful predictions on randomized data with a similar variant distribution. The software does, however, issue meaningful predictions based on the non-randomized data.

In our analysis, we focused specifically on non-synonymous variants, as these variants are expected to directly affect protein function. Despite this, all the SNPs associated with ASD in this study are considered benign. This observation is consistent with findings from other studies, which suggest that the influence of SNPs on complex traits such as ASD often depends on their collective, rather than individual, effects. Moreover, similarly to GWAS studies, these SNPs may be in linkage disequilibrium (LD) with causal variants.

Ultimately, while non-synonymous and frameshift SNPs are expected to have functional consequences, the combinatorial effect of multiple SNPs, as well as their interaction with regulatory variants, may have a more profound impact on gene expression and ASD pathogenesis. Further functional studies are necessary to explore these combinatorial effects and to better understand how these variants influence neurodevelopmental pathways in ASD.

The 177 genes identified in this study exhibited significant overlap with the SFARI Gene database, which is focused on genes associated with ASD risk through analysis of de novo and rare variants. The convergence of genes identified by rare variants with our results supports the validity of our approach and strengthens the evidence for the involvement of these genes in ASD pathogenesis. In addition, identification of ontologies already associated with ASD, such as neuron development processes, provides further support for our findings.

At the connectivity analysis level, we observed low connectivity group of 144 genes and high connectivity group of 33 genes, and the presence of 3 key hub genes, MYLK3, CLIP1 and TNC. The low connectivity group showed enrichment mainly to abnormality of the eye and abnormal eye physiology. While the high connectivity group showed enrichment to immune system process and ASD related behavioral phenotypes. The hub genes MYLK3 and TNC and their co-occurring genes were enriched for multiple neuronal processes, while the CLIP1 did not show such enrichment. The different enrichments according to connectivity may indicate different genetic mechanisms influencing the phenotypes. However, the meaning of gene connectivity in this study should be further investigated to understand whether it represents a genetic mechanism.

One limitation of this study was the lack of detailed phenotypic information. While our analysis identified several SNPs associated with ASD, the disorder’s phenotypic heterogeneity complicates the interpretation of these findings. Despite this, the analysis revealed a novel association of specific abnormality of the eye genes and ASD. In spite of the fact that atypical eye gaze responses are common in ASD, and show high heritability, this phenotype has never been studied either in the context of rare or common variants. The association of 49 genes related to eye abnormality with ASD highlights the advantage of studying specific phenotypes rather than global umbrella binary phenotypes in genomic studies, which will be valuable for identifying novel variants increasing ASD risk. Identifying NOTCH1 signaling and low voltage-gated calcium channels in the context of eye physiology may pave the way to medical ASD treatment. Future research should aim to validate our findings in independent cohorts and to further investigate the functional consequences of the identified variants.

Finally, our novel findings on ASD genetic underpinnings highlight the power of machine learning approaches to identify common genetic variants associated with complex disorders.

## Methods

### Datasets

Exome sequencing data from the Autism Sequencing Consortium(21) (ASC, phs000298.v4.p3) a cohort of 8,061 individuals—3,585 diagnosed with autism and 4,476 controls. For validation we used controls from the SCZ Population-Based Case–Control Exome Sequencing study (phs000473.v2.p2), comprising 5890 participants.

### Variant calling and annotation

Reads were mapped to the UCSC Human reference genome (GRCh37/hg19) using BWA(22) and variants were identified using GATK(23). The SNP annotation of the variants was performed using SnpEff(24) with dbSNP as the reference database. SnpSift(25) was used to annotate SNP protein-affecting variants, which were selectedb for further analysis.

### Variant matrix creation

For each gene in the dataset, we calculated the frequency of each SNP among all the subjects in the dataset. Then, for each subject, we assigned the most common variant among the SNPs that was identified for this subject in each of the genes. For the ASD dataset, this process resulted in a matrix of 8061 rows (number of individuals in the ASD study) and 18497 columns (18497 genes harboring 655780 variants).

### Machine learning

We used WizWhy(19), a machine learning software program designed to associate independent variables (e.g., genetic variants) with the dependent variable (e.g., phenotype). WizWhy works by revealing all the if–then rules that associate the independent variables with the dependent variable and meet user-determined thresholds for rule confidence (minimum probability inversely proportional to the frequencies of the dependent variable) and support (minimum number of cases = 40). Given the dataset’s extensive number of variants and relatively limited number of records, we constrained the search to rules with two conditions. We performed 5 cross validations by randomly dividing the dataset into training and test sets, with rules being discovered in the training set, and then using these rules, generating predictions for the test set. A key advantage of using WizWhy is that the rules provide explanations for the predictions and can be used to build hypotheses that explain the data.

### Validation

To validate the rules, Fisher exact test was performed for each variant pair discovered by WizWhy with the null hypothesis that its occurrence in the SCZ control is equal or higher than expected. Rules with FDR smaller than 0.05 were selected for further analysis.

### Enrichment analysis

177 unique genes were analyzed to assess gene enrichment. Gene enrichment analysis was performed by ShinyGO(26) v 0.82 using a hypergeometric test.

### Connectivity analysis

Co-occurrences were determined by counting the number of appearances of each gene in the rules. The density of connectivity was generated with R’s ggplot2.

### Network construction and analysis

To analyze the relationships between genes identified in association rules, we constructed an undirected network where nodes represent genes and edges represent co-occurrence in WizWhy association rules. Each rule contained gene modifications that were correlated with disease occurrence above a specified confidence threshold. The final WizWhy-derived association network contained 177 genes (nodes) and 533 edges. Network parameters analysis and visualization were performed using Cytoscape v.3.8.2.

To identify biologically meaningful genes within this network, we performed topological analysis focusing on hub identification.

We analyzed the network using four complementary topology metrics from the Cytoscape plugin cytoHubba: degree, maximal clique centrality (MCC), betweenness, and closeness. For each metric, two thresholds were applied to identify genes with significant network importance: a statistical threshold set at μ + 2σ (μ = mean, σ = standard deviation of scores) and a percentile-based threshold (top 5% of ranked genes). To ensure robustness, we defined hub genes as those surpassing these thresholds in at least two metrics, thus capturing nodes of high topological significance.

### Statistics

Statistical analyses were performed using R. The hypergeometric test was performed to determine the significance of the overlap between genes in our analysis and SFARI gene database.

## Conflict of interest

The authors declare the following competing interests:

Alona Rabner, Assaf Avrahami, Abraham Meidan, and Ilya Vorobyov are employees of WizSoft.

WizSoft also partially funded the work of Guy Horev.

Abraham Meidan is a co-owner of WizSoft.

